# Melanin concentrating hormone and orexin shape social affective behavior via action in the insular cortex of rat

**DOI:** 10.1101/2023.03.09.531898

**Authors:** Lucas Barretto-de-Souza, Shemar A. Joseph, Francesca M. Lynch, Alexandra J. Ng, John P. Chrsitianson

## Abstract

**Rationale:** In a social context, individuals are able to detect external information from others and coordinate behavioral responses according to the situation, a phenomenon called social decision-making. Social decision-making is multifaceted, influenced by emotional and motivational factors like stress, sickness and hunger. However, the neurobiological basis for motivational state competition and interaction are not well known.

**Objective:** We investigated possible neural mechanisms through which internal states could shape social behavior in a social affective preference (SAP) test. In the SAP test, experimental rats given a choice to interact with naïve or stressed conspecifics exhibit an age-dependent preference to interact with stressed juvenile conspecifics, but avoid stressed adult conspecifics. First, we assessed the effect of hunger on SAP behavior. Behavior in the SAP test requires the insular cortex, which receives input from the hunger-related peptides melanin-concentrating hormone (MCH) and orexin neurons of the lateral hypothalamus (LH). This study aimed to evaluate the role of LH and insular MCH and orexin in SAP test.

**Methods:** SAP tests were conducted in rats that were sated, food deprived or allowed 1 h of access to food after 14 h of deprivation (relieved condition). Separate cohorts of sated rats received cannula implants for microinjection of drugs to inhibit the LH or to block or stimulate MCH or orexin receptors in the insula prior to SAP tests or social interaction tests.

**Results:** Food and water deprivation prior to SAP tests with juvenile rats caused a shift in preference away from the stressed rat toward the naïve juveniles. Pharmacological inhibition of LH with muscimol (100 ng/side) abolished the preference for the juvenile stressed conspecific, as well as the preference for the adult naïve conspecific. The blockade of MCHr1 or orexin receptors in the insular cortex with SNAP94847 (50µM) or TCS1102 (1µM), respectively, also abolished the preference for the stressed juvenile conspecific, but only the antagonism of orexin receptors was able to abolish the preference for the adult naïve conspecific. Microinjection of increasing doses (50 or 500 nM) of MCH or orexin-A in the insular cortex increased the interaction time in the one-on-one social interactions test with juvenile conspecifics, however only the microinjection of orexin-A increased the interaction time with adult naïve conspecifics.

**Conclusions:** Taken together, these results suggest that lateral hypothalamus peptides shape the direction of social approach or avoidance via actions MCH and orexin neurotransmission in the insular cortex.

## INTRODUCTION

The ability to generate appropriate behaviors in different social contexts is crucial for survival and adaptation of social animals (Insel and Fernald 2004). This dynamic process involves successive prediction and evaluation of the motivations and emotions of others (Ranote et al. 2004; Rendon et al. 2015). Behavioral responses to others are also influenced by the environment and internal states such as stress, sickness or hunger (Sandi and Haller 2015; Toyoshima et al. 2021, 2022; Devlin et al. 2022; Smith and Grueter 2022). In rodents, the interaction of these factors results in a range of prosocial and avoidance behaviors directed toward distressed conspecifics (Meyza et al. 2017). Behaviors observed in rodent social interactions are thought to be the product of the social decision-making network (SDMN) (O’Connell and Hofmann 2011), a highly conserved system of brain regions comprised of the mesocorticolimbic reward pathway and the “social brain network” (Newman 1999).

A region highly connected to the SDMN is the insular cortex (reviewed by Rogers-Carter and Christianson, 2019). Insula is a site of multisensory integration implicated in a wide range of cognitive domains (Kurth et al. 2010; Gogolla 2017). With regard to social behavior, the insula is necessary for both prosocial and avoidant responses to socioemotional stimuli. In the social affective preference (SAP) test, experimental rats exhibit age-specific responses to adult or juvenile rats that were exposed to stress. Specifically, when presented with a pair of adult conspecifics, one stressed and one naïve to stress, experimental rats avoid interactions with the stressed conspecifics. In contrast, when presented with a pair of prepubescent juveniles, experimental rats spend more time investigating the stressed conspecific. Treatments that inhibit the activity of the insula during SAP tests interfere with this pattern (Rogers-Carter et al. 2018, 2019; Rieger et al. 2022). The insula appears to be an integral circuit component linking social and situational factors to inform social decisions that, via interaction with the SDMN, can shape the circuit activity pattern to favor the approach or avoidance behavior strategies. Consistently, inhibition of affective input from the basolateral amygdala to the insula (Djerdjaj et al. 2022) or inhibition of insula projections to the nucleus accumbens both interfere with the approach preference for stressed juvenile animals (Rogers-Carter et al. 2019).

As noted above, social interactions are influenced by a range of internal factors and understanding the biological mechanisms by which competing hunger and social motivations are resolved is fundamentally interesting. Importantly, hunger or direct stimulation of hunger associated neural circuits can reduce sociability (Burnett et al. 2016, 2019) which was suggested to be the consequence of the need to find food taking priority over needs for social interaction in the brain structures responsible for decision making (Sutton and Krashes 2020). To determine if hunger might influence social interactions in the SAP test, we conducted a series of three SAP tests in male rats that were sated, with free access to food and water; hungry, after 14 hours of food and water deprivation; or relieved, after 13 hours of food deprivation with 1 hour of free access prior to test. As usual, when tested in a sated state, rats spent more time investigating the stressed juvenile, but in either food deprived state, rats spent more time investigating the naïve juvenile (Figure 1). These observations and the large body of prior research implicating the insula in several aspects of appetitive and consummatory behavior (Yiannakas and Rosenblum 2017; Livneh and Andermann 2021) suggested to us that hunger signals could influence social decision making, in part, by action in the insular cortex.

**Figure 1.**
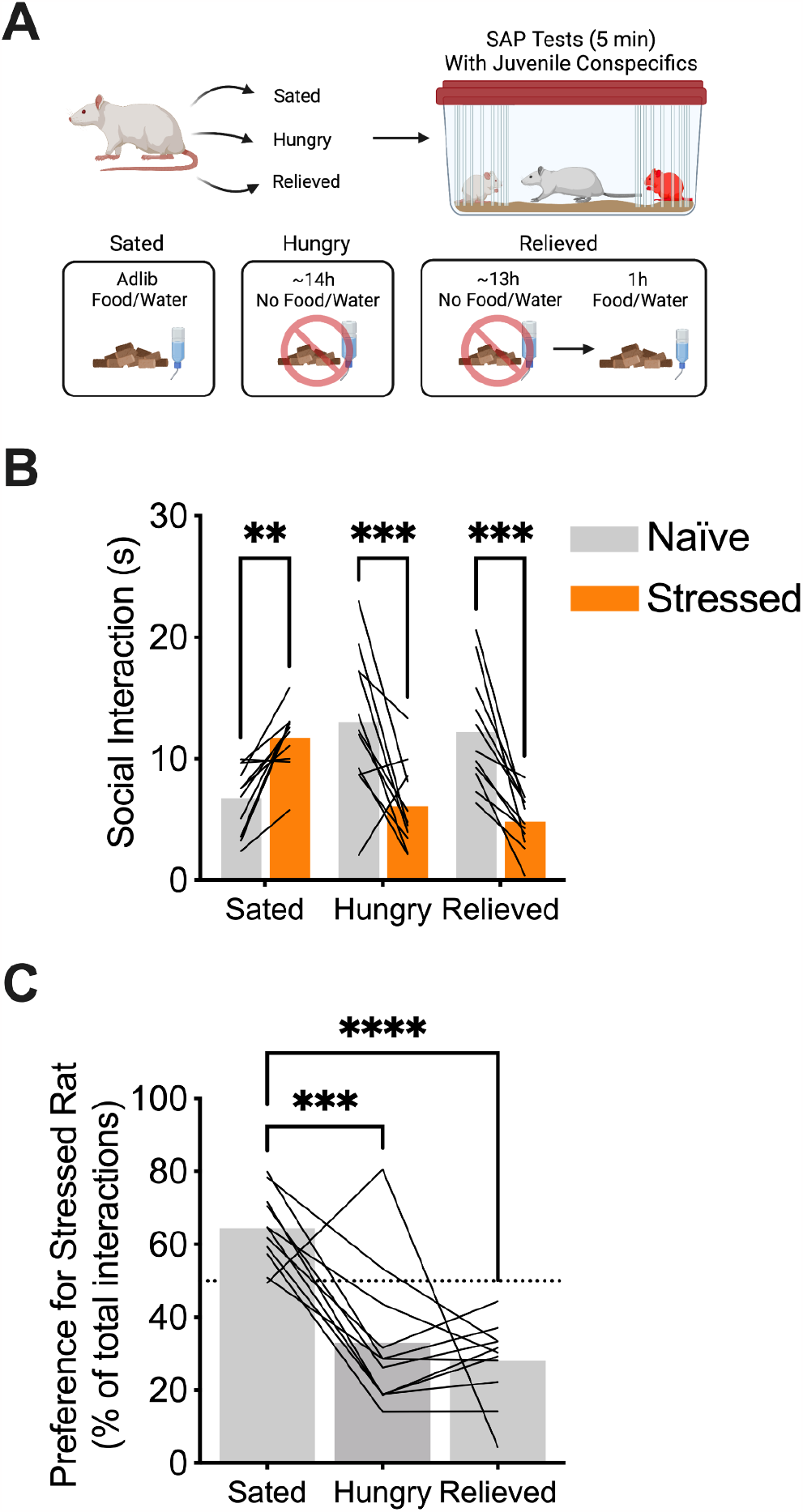
Hunger interferes with social affective behavior. **A.** Schematic figure of the experimental protocol, rats (n=11) under sated, hungry, or relieved conditions underwent SAP test with juvenile conspecific. **B.** Mean (with individual replicates) time spent interacting with each conspecific during the 5 min SAP test. Sated rats spent more time interacting with the stressed juvenile conspecific, but hungry and relieved rats spent more time interacting with the naïve juvenile conspecific. **C.** Data from B converted to preference for stressed conspecific in percentage of total interactions (mean with individual replicates). Sated rats presented a preference (>50% time interaction) for the stressed conspecific, however either hungry condition reduced significantly this preference. Panel A created with BioRender.com. **p<0.01, ***p<0.001, ****p<0.0001.

Acute hunger drives exploratory and consummatory behaviors through well defined hypothalamic circuits. Briefly, as caloric deficits increase, Agouti-related peptide (AgRP) neurons in the arcuate nucleus ramp up activity in axons that project to the lateral hypothalamus (LH) (Andermann and Lowell 2017). The LH is a complex diencephalic region (Saper et al. 1979) involved in the control of behaviors related to arousal, feeding, energy balance, stress, reward and motivated behavior (Bonnavion et al. 2016; Petrovich 2018). Input from AgRP neurons evokes activity in LH neurons containing the neuropeptides melanin concentrating hormone (MCH) and orexin/hypocretin which are among a diverse group of LH neurons that project widely through the brain (Swanson et al. 2005; Hahn 2010), including to the insular cortex which contains MCHr1 and both orexin 1 and 2 receptors (Hervieu et al. 2000; Li and de Lecea 2020; Mitsukawa and Kimura 2022). MCH and orexin systems are vital to feeding and energy balance, however they are also related to sleep, stress, arousal, and other motivated behaviors (Sakurai et al. 1998; Huang et al. 2007; Matsuki et al. 2009; Bonnavion and de Lecea 2010; Kaplan et al. 2022). A few recent reports manipulations of the LH itself, MCH, or orexin systems suggest they contribute to various aspects of social behavior (Blouin et al. 2013; Abbas et al. 2015; Nieh et al. 2016; Noritake et al. 2020; Sanathara et al. 2021), leading us to hypothesize that these may be components of the neural circuit through which internal motivational states are integrated with social affective information to shape social behavior. Although orexin receptors in the insula contribute to feeding behavior (Hagar et al. 2017), the role of these LH neuropeptides in the insula on social behavior is unknown. Here we report on a set of experiments in which we evaluate the role of LH itself and MCH and orexin neurotransmission in the insula in the social approach/avoidance behavior in the SAP tests.

## Materials and Methods

### Subjects

Male Sprague Dawley rats were obtained at either post-natal day PN50, PN22 or PN42 from Charles River Laboratories. All rats were acclimated to the vivarium in the Boston College Animal Care facility for a minimum of 7 d before surgery or behavioral testing, resulting in groups of test adult rats at PN60–PN80, juvenile conspecifics at PN30 and adult, post pubescent conspecifics at PN50 at the time of testing. Subjects were housed in groups of 2 or 3, maintained on a 12 h light/dark cycle, and provided food and water ad libitum. Behavioral testing occurred within the first 4 h of the light cycle. All procedures were approved by the Boston College Institution Animal Care and Use Committee and adhered to the National Institutes of Health’s Guide for the Care and Use of Laboratory Animals.

### Summary of Experiments

#### Influence of food deprivation or relief on SAP behavior

Over 7 days, rats (N=11) were given SAP tests with juvenile conspecifics under sated, hungry, or relieved conditions (See Figure 1). On Days 1 and 2 rats were habituated to the SAP arena and on Days 3, 5 and 7 SAP tests were performed. Rats were subdivided into 3 groups to allow for counterbalancing of treatment order. Food restriction entailed removal of food and water from the homecage about 14 h prior to SAP testing. For the relief condition, food and water were replaced on the cage for 1 h prior to SAP. Sated rats were allowed free access to food. Following SAP tests, regardless of condition, rats had free access to food and water for ∼ 48 h. Test order was counterbalanced in a latin square design and no order effects were evident.

#### Inhibition of the LH during SAP tests

Rats were implanted with cannula guides for microinjection of muscimol to the LH. After recovery, SAP tests were performed over 5 days; experimental rats were tested with either juvenile or adult conspecifics (See Figure 2). Days 1 and 2 were habituation days and on Day 3 and 5, SAP tests 1 h after injection of muscimol or vehicle. No testing occurred on Day 4 to ensure complete wash out of the drug.

**Figure 2.**
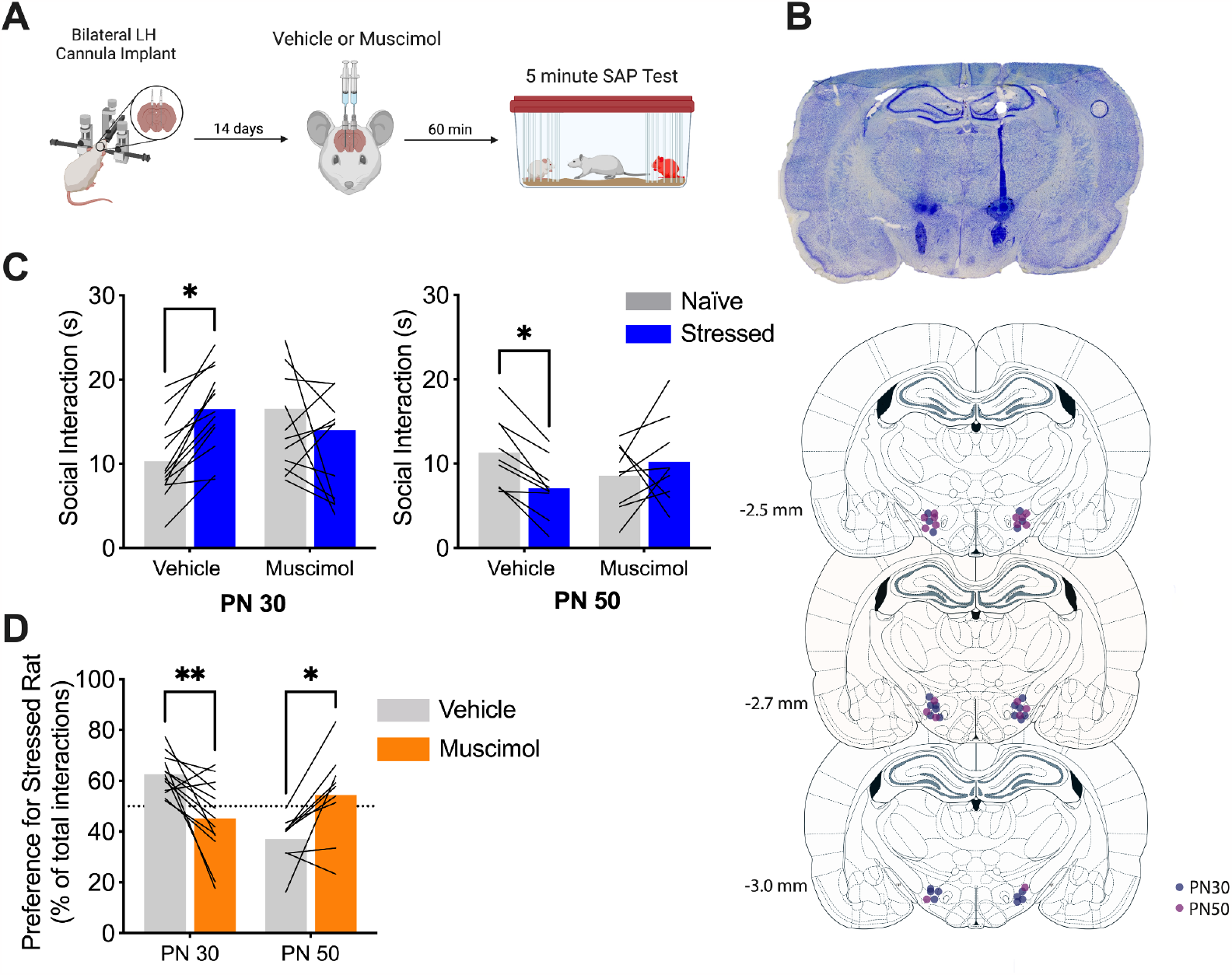
Pharmacological inhibition of the LH interfered with social affective behavior in the SAP test. **A.** Schematic representation of the experimental protocol. Rats had bilateral cannula implanted in the LH and after 2 weeks of recovery rats received microinjections of vehicle or muscimol (100 ng/side) into the LH and 60 min later underwent SAP test with juvenile or adult conspecifics. **B.** Representative image of bilateral cannula tract in the LH and maps of cannula placements of the LH microinjections of rats tested with juvenile (dark purple, n = 13) or adult (light purple, n = 9) conspecifics. **C.** Mean (with individual replicates) time spent interacting with each conspecific during the 5 minutes of SAP tests with juveniles (left, PN 30) or adults (right, PN 50). Test rats spent more time interacting with the stressed juvenile conspecific, this behavior was not evident after administration of muscimol. In SAP tests with adults, rats spent more time interacting with the naïve adult conspecific, which was not evident after administration of muscimol. **D.** Data from C were converted to preference for stressed conspecific which is the percentage of interaction with the stressed conspecific out of the total interactions with both naïve and stressed conspecifics (mean and individual replicates shown). Rats tested with juvenile conspecific presented a preference (>50% time interaction) for the stressed conspecific and those tested with adults preferred the naïve conspecific. Treatment with muscimol reduced the preference for stressed juveniles and naïve adults. *p<0.05, **p<0.01. Panel A created with BioRender.com.

#### Insula MCH and orexin in social interaction

Rats were implanted with cannula guides to the posterior insula, the region previously identified as necessary for SAP behaviors. After recovery, SAP tests were performed with either adult or juvenile conspecifics over 4 days. Days 1 and 2 were habituation days and on Days 3 and 4 SAP tests were conducted 30-40 min after infusion of MCH receptor 1 (MCHr1) antagonist or dual orexin 1/orexin 2 receptor antagonist in counterbalanced order. Separate cohorts of rats were used for each SAP age and drug treatment. To determine if insular MCH or orexin would affect sociability in the absence of stress, in the week after SAP testing, we conducted 5 min social interaction tests with either juvenile or adult conspecifics 30-40 min after infusion of MCH or orexin-A at 2 different doses.

### Social affective preference test

Initially described by Rogers-Carter, Varela et al., (2018), the Social Affective Preference (SAP) test allows observation of test rat behavior when presented simultaneously with 2 conspecifics, one of them stressed and the other naïve to any treatment. The test takes place in a plastic arena (76.2 × 20.3 × 17.8, cm; L × W × H) lined with beta chip bedding in which the test rat can move freely. At the extremities, the conspecific animals are positioned inside an acrylic box (18 × 21 × 10 cm; L × W × H) with rods spaced 1 cm apart to allow interaction with the test rat. The experimental protocol consists of 2 habituation days followed by 2 test days. On day 1 the test rats are habituated to the arena with the empty conspecific box for 15 minutes and stay in the experiment room in individual cages for 60 min. On day 2, after 60 minutes of habituation in their respective individual cages, test animals are presented with 2 naïve conspecifics for 5 min in the testing arena. For food deprivation/relief experiments, on days 3, 5 and 7 the test animals underwent SAP tests with juvenile conspecifics under either sated, hungry or relieved conditions. Treatment order was counterbalanced across days (within-subjects design). For pharmacological experiments, on days 3 and 5 (inhibition of the LH during SAP tests experiment) or on days 3 and 4 (insula MCH and orexin in social interaction experiments), the test animals received microinjection of vehicle or drug, counterbalanced between days (within-subjects design), and underwent a simultaneous interaction with a stressed and a naïve to any treatment unfamiliar conspecifics for 5 minutes. Conspecific stress entailed exposure to 2 footshocks (5 s, 1 mA, inter-shock interval of 50 s (in a separate sound and odor isolated room) immediately before placement in the SAP test arena. Interactions always occurred with a novel conspecific. Interaction time with conspecifics are scored by a trained observer and all tests were recorded on digital video to establish inter-rater-reliability.

### Lateral hypothalamus and insular cortex cannula placement

Cannulas (26-gauge, Plastics One, Roanoke, VA) were inserted inserted bilaterally into the LH (from bregma: AP: –2.7 mm, ML: ±2 mm, DV: –7.8 mm from skull surface) or into the posterior insula (from bregma: AP: –1.5 mm, ML: ±6.2 mm, DV: –6.8 mm from skull surface) under inhaled isoflurane anesthesia (2–5% v/v in O2) in a stereotaxic frame. After surgery, rats received meloxicam (1 mg/kg, Eloxiject; Henry Schein) and the antibiotic penicillin (12,000 units, Combi-pen; Henry Schein) and were allowed to recover undisturbed for 2 weeks before behavioral testing.

### Pharmacological manipulations

All drugs were obtained from Tocris Bioscience. For pharmacological inhibition of LH cell bodies, test rats received LH injections of 100 ng/side of the selective GABA_A_ agonist muscimol dissolved in sterile 0.9% saline 60 min before SAP testing. This dose was selected based on prior experience in our laboratory (Rogers-Carter et al. 2018). For the pharmacological inhibition of MCHr1 or orexin receptors, experimental rats received, respectively, the selective MCHr1 antagonist SNAP94847 (50µM) or the dual orexin receptor antagonist TCS1102 (1µM). Both antagonists were first dissolved in DMSO and diluted to final concentration in saline. For the activation of the MCHr1 and orexin receptors, the endogenous agonists MCH and orexin-A, respectively, both at 50 or 500nM, dissolved in saline. All the injections had a volume of 0.5 μl/side. Doses were selected based on the Ki values provided by manufacturers and used at doses comparable or less concentrated than previously reported for SNAP94847 (Hsiao et al. 2012; Rodrigues et al. 2022) or TCS1102 (Kokare et al. 2006; Sabetghadam et al. 2018; Li et al. 2021; Liu et al. 2022).

### Social Exploration

One-on-one social interaction tests were performed on days 3, 4 and 5 after the end of the SAP test to determine if MCH or orexin-A administration to the insula was sufficient to alter social behavior in the absence of stress. On each day, the experimental animals received microinjection into the insula of vehicle or agonist and 45 minutes later a conspecific naïve to any treatment was introduced into their home cage for 5 minutes. Three tests allowed for a within-subjects dose-response experiment with each rat tested after vehicle, 50 and 500μM doses of MCH or orexin-A. Drug order was counterbalanced using a latin square design. During the entire test period, interaction times with conspecifics were scored by a trained observer and all of the tests were recorded in a digital video.

### Tissue collection and Histology

After the last day of the experimentation, test animals were overdosed with tribromoethanol (Fisher Scientific). Brains were removed and stored in a 10% paraformaldehyde solution at 4°C for at least 48h and then transferred to 30% sucrose for at least 48h. Coronal sections (40μm) of the LH or insula were mounted on a gelatin-coated slide using a freezing cryostat (Leica) and subsequently stained with cresyl-violet for verification of cannula placement.

### Data analysis

Data from animals with incorrect placement were removed from the analysis. In our initial report about SAP test behavior, we observed a normal distribution of preferences with a fraction of rats sometimes displaying avoidance of stressed juveniles, or approach to stressed adults (Rogers-Carter et al. 2018). In our experience, experiments involving surgery and injections can increase the frequency of subjects exhibiting unusual behavior, specifically in SAP tests with adults where social interaction levels are somewhat lower than with juveniles. As we have previously described (Rogers-Carter et al. 2019), a few experimental subjects were excluded from analysis of adult SAP tests with LH muscimol and insula orexin antagonist experiments because the rats exhibited fewer than 5 seconds of total interaction, or exhibited a preference for the stressed adult conspecific greater than 50%. These instances are noted in the corresponding results sections. Social interaction time during the SAP test was analyzed using repeated measures two-way ANOVA with conspecific stress and drug treatment (vehicle or antagonist) or hunger state (sated, hungry or relieved) as within-subjects variables. The preference for the stressed conspecific during SAP test was calculated as a percentage of time interacting with the stressed conspecifics relative to the total time spent interacting both conspecifics and was analyzed using repeated measures two-way ANOVA with conspecific age and drug treatment (vehicle or antagonist) as within-subjects variables or using repeated one-way ANOVA with hunger state as within-subjects variable. Social interaction time during the one-on-one social interaction test was analyzed using repeated measures two-way ANOVA with conspecific age and drug treatment (vehicle or agonist) as within-subjects variables. A significant effect was considered when p < 0.05 and post-hoc analysis consisted of Šidák’s multiple comparison tests.

## RESULTS

### Acute food and water deprivation reversed preference for stressed juvenile conspecifics

To test whether hunger would influence the approach behavior toward stressed juvenile conspecifics, test rats underwent SAP tests with juvenile conspecifics under sated, hungry, or relieved conditions (Figure 1A). As expected, sated rats displayed preference for the stressed juveniles, but hungry and relieved rats avoided the stressed juvenile (Figure 2B). A two-way ANOVA indicated a conspecific stress main effect (F(1, 10) = 11.48; p = 0.0069) and a hunger by conspecific stress interaction (F(2, 20) = 22.60; p < 0.0001). Post-hoc analysis indicated that animals under sated condition preferred to interact with the stressed juvenile conspecific (p = 0.0091). However, animals under either hungry (p = 0.0004) or relieved (p = 0.0002) conditions spent more time interacting with the naïve juvenile conspecific. The preference of stressed conspecific was calculated as percentage and analyzed with a one-way repeated measures ANOVA which revealed a main effect of hunger state (F(2, 20) = 19.61; p = 0.0006). Post-hoc analysis indicates an opposite preference for the stressed juvenile conspecific in either hungry (p = 0.0035) and relieved (p < 0.0001) conditions when compared with sated condition (Figure 1C).

### Pharmacological inactivation of LH abolished both preference for stressed juvenile and naïve adult conspecifics

To test whether the LH is necessary for social affective behavior during SAP test, rats underwent SAP test with either juvenile (n = 13) or adult conspecifics (n = 9) after 60 minutes of vehicle or muscimol (selective GABA_A_ agonist) (100 ng/side) microinjection in the LH (Figure 2A,B,C). Four rats in the adult condition were excluded from analysis due to unusual SAP behavior after vehicle injections. Analysis of juvenile SAP tests evidenced a significant conspecific stress by drug interaction (F(1, 12) = 8.747; p=0.0120) in rats tested with juvenile conspecific and post hoc tests confirmed that after vehicle injections rats have a preference to explore the stressed conspecifics (p = 0.0232), however this was not evident after the pharmacological inactivation of LH (p = 0.4364) (Figure 2D). When tested with adult conspecifics the analysis indicated a significant conspecific stress by drug treatment interaction (F (1, 8) = 11.62; p = 0.0092), the post-hoc analyses indicated a preference to the adult naïve conspecific (p=0.0168) under vehicle condition, however muscimol abolished this behavior (p=0.3830) (Figure 2E). The preference for stressed conspecific was calculated as percentage, data analysis indicated a significant conspecific age by drug interaction (F (1, 20) = 22.52; p=0.0001), and LH inhibition abolished the preference for stressed juvenile (p=0.0027), as well the avoidance for stressed adult conspecific (p=0.0117) (Figure 2F).

### Melanin concentrating hormone, orexin, and social affective behavior

To determine if the LH neuropeptides MCH and orexin would influence social behavior by action in the insula, experimental rats received bilateral cannula in the posterior insula and then underwent the SAP tests with receptor antagonist or vehicle injections. We previously found other neuromodulator receptors including oxytocin and corticotropin releasing factor 1 receptors to be necessary for SAP behavior and in each case, simply administering the agonist for these receptors was sufficient to mimic the effect of conspecific stress on social interaction. For example, oxytocin administered to insula increases social interactions with naïve juveniles but reduces interaction with naïve adults, suggesting that this form of neurotransmission is sufficient to mediate the behaviors observed in the SAP test (Rogers-Carter et al. 2018). To determine if MCH or orexin are sufficient to influence social interaction, 3 days after the SAP tests rats received a series of 3 one-on-one social interaction tests with either adult or juvenile conspecifics after vehicle or either MCH or orexin-A infusions (Figure 3).

**Figure 3.**
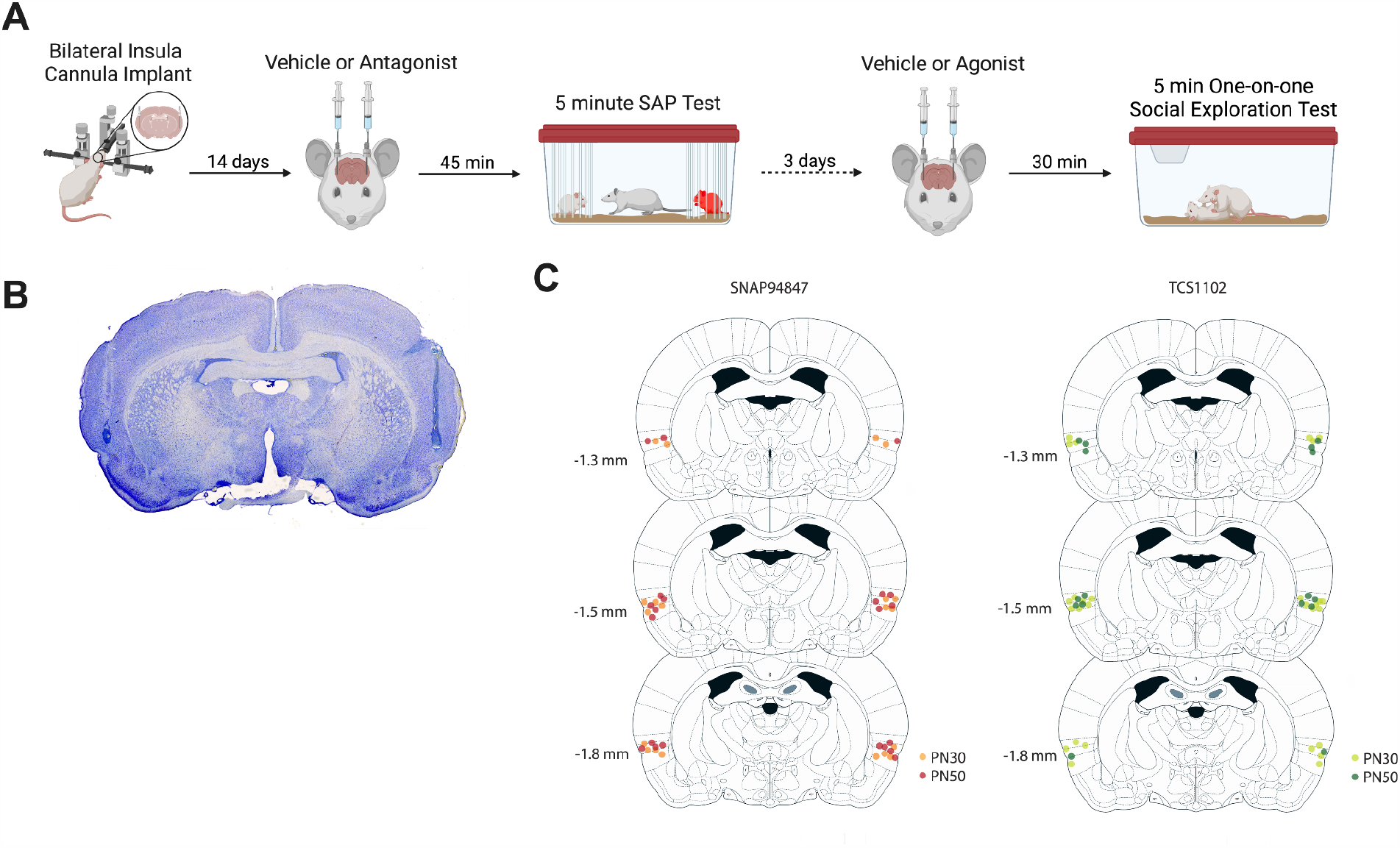
Overview of MCH and orexin experiments. **A.** Schematic figure of the experimental protocol. Cannula were implanted in the insula and after 2 weeks of recovery rats received microinjections of vehicle or either MCHr1 antagonist (SNAP94847) or dual orexin receptor antagonist (TCS1102) into the insula and 45 minutes later underwent SAP tests with juvenile or adult conspecifics. Separate groups of rats were used for adult and juvenile tests, and each group was only tested with one set of drugs targeting either MCH or orexin receptors. SAP tests were repeated the next day with the opposite drug treatment in a within subjects design; drug order was counterbalanced. After 3 days without treatments, rats underwent a series of 3 one-on-one social interaction tests after either vehicle or one of 2 doses of MCH or orexin-A to determine if administration of the ligand would alter sociability itself. Rats were given 1 social interaction test per day, treatment order was counterbalanced, and rats were tested with the same aged conspecifics as in the prior SAP tests. Social interaction tests were conducted over 5 minutes with a test rat and an unfamiliar, naïve juvenile or adult conspecific. **B.** Representative image of bilateral cannula tract in the insula. **C.** Placement maps of the insula microinjections of SNAP94847 in rats tested with juvenile (n = 10) or adult (n = 11) conspecifics (left) and of TCS1102 in rats tested with juvenile (n = 14) or adult (n = 8) conspecifics (right).

To evaluate the role of insular MCH neurotransmission on social affective behavior, SAP tests were conducted in rats that received either vehicle or the selective MCHr1 antagonist SNAP94847 (Figure 4). Analysis of juvenile SAP tests (n = 10) indicated a significant conspecific stress by drug treatment interaction (F(1, 9) = 12.56, p = 0.0063) and post hoc tests indicated a preference for the stressed juvenile conspecific after the treatment with vehicle (p = 0.0070), but this was not present after administration of the SNAP94847 (p = 0.5143) (Figure 4A). Data from rats tested with adult conspecifics (n = 11) presented a main effect of conspecific stress (F(1, 10) = 5.1619; p = 0.0393). Although a stress by drug interaction only approached significance (F(1, 10) = 4.169; p = 0.0684), post-hoc analysis indicated that rats treated with vehicle had a preference for naïve adult conspecific (p = 0.0087) but the time spent interacting with naïve and stress conspecifics did not differ after administration of SNAP94847 (p = 0.7025) (Figure 3B). The preference of stressed conspecific was calculated as percentage and analysis indicated a main effect of conspecific age (F(1, 19) = 15.11; p = 0.0010) and a significant conspecific age by drug treatment interaction (F(1, 19) = 14.98; p = 0.0010), the MCHr1 inhibition prior the SAP test abolished the preference for stressed juvenile (p = 0.0021), but not the avoidance for adult stressed conspecific (p = 0.2496) (Figure 4C).

**Figure 4.**
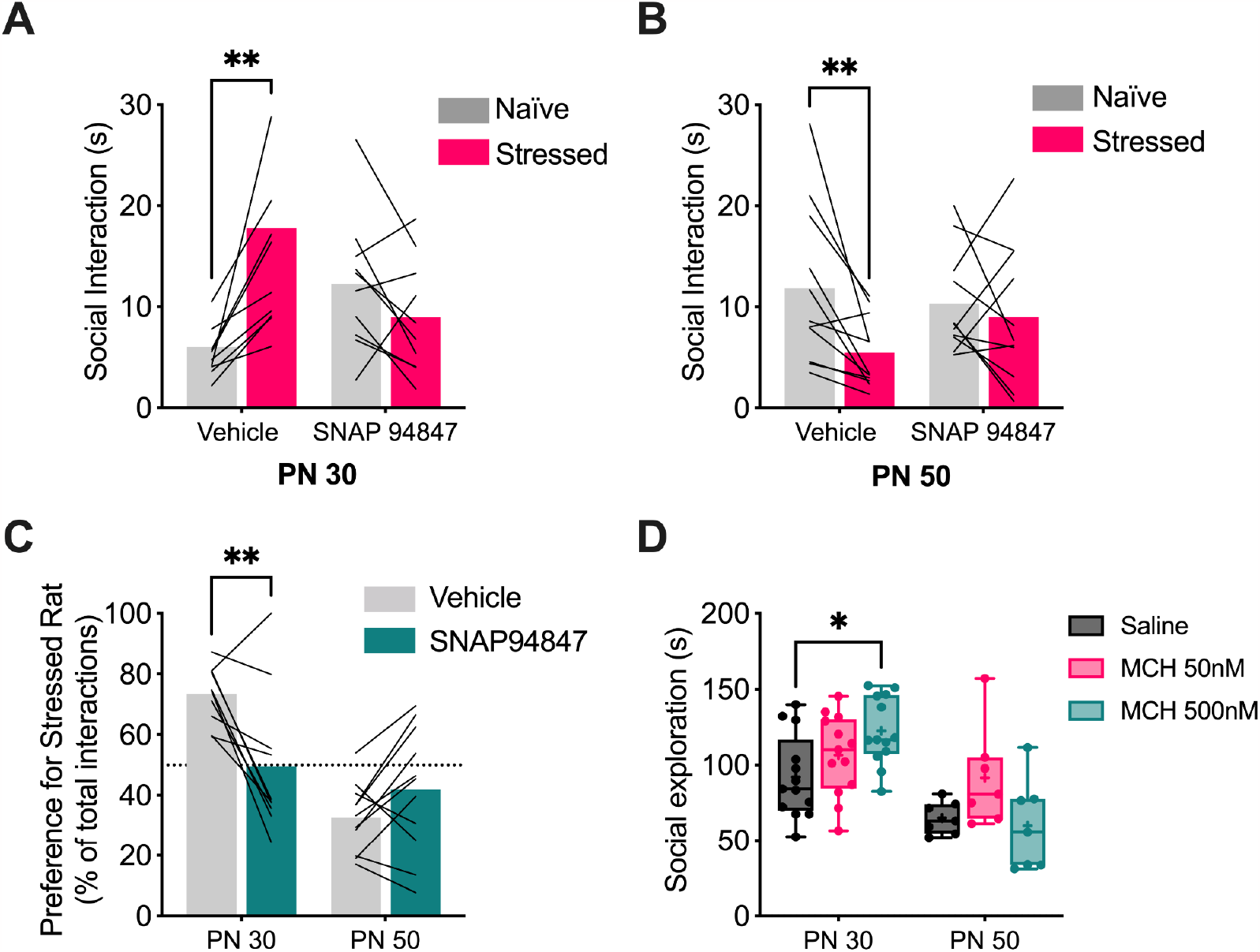
MCH neurotransmission affects social approach behavior to juveniles but not avoidance of adults. Juvenile and adult SAP tests were conducted 45 min after administration of the vehicle or the MCHr1 antagonist SNAP94847 (50µM) to the insula. **A.** Mean (with individual replicates) time spent interacting with juvenile conspecifics during the SAP test. Rats spent more time interacting with the stressed juvenile conspecific which was not evident after injection of SNAP94847. **B.** Mean (with individual replicates) time spent interacting with adult conspecifics during the SAP test. Test rats spent more time interacting with the naïve adult conspecific, which was not evident after infusion of administration of SNAP94847. **C.** Preference for the stressed conspecific expressed as percentage of total interaction time (mean with individual replicates). Rats tested with juvenile conspecifics presented a preference for the stressed conspecific which was significantly reduced by SNAP94847. In SAP tests with adults, SNAP9447 did not consistently alter preference. **D.** Time spent exploring naïve juvenile and adult conspecifics in a 5 minute social interaction test (box depicts median and first and third quartile, whiskers indicate maximum and minimum, and the mean is indicated by a +) 30 min after insula injections of vehicle or MCH (50 and 500 nM). MCH increased time spent exploring naïve juveniles but did not influence interactions with adult conspecifics. *p<0.05, **p<0.01.

On day 3, 4 and 5 after the end of the SAP test, test rats received microinjection of vehicle or increasing doses of MCH (50 and 500 nM) in the insula and were submitted to one-on-one social interaction test with naïve juvenile or adult conspecific after 30 minutes. MCH appeared to selectively increase social interactions directed toward juveniles (Figure 4D). Analyses revealed a main effect of conspecific age (F(1, 18) = 19.68; p = 0.0003), and a significant conspecific age by drug treatment interaction (F(2, 36) = 4.747; p = 0.0148). The post-hoc test indicated that only rats tested with naïve juvenile conspecifics and treated with the highest dose of MCH (500nM) in the insula showed an increase in interaction time during the one-on-one social exploration test (p = 0.0240) (Figure 4D).

To test whether orexin action in the insula has an influence on SAP behavior and one-on-one social exploration test rats were submitted to the same protocol described above, but received microinjection of vehicle or TCS1102 (1µM) 45 minutes before SAP tests (Figure 5). Analysis of SAP tests with juveniles (n = 14) revealed a significant conspecific stress by drug treatment interaction (F(1, 13) = 13.07; p = 0.0031). Post-hoc test indicated a preference for the stressed juvenile conspecific after the treatment with vehicle (p = 0.0357), but this was not present after TCS1102 (p = 0.0622) (Figure 5A). Analysis of SAP tests with adult conspecifics (n = 8, 4 rats were excluded from analysis because of unusual behavior under vehicle, Figure 5B) presented a significant conspecific stress by drug treatment interaction (F(1, 7) = 8.909; p = 0.0204). Post-hoc analysis indicated that rats treated with vehicle spent more time interacting with the naïve adult conspecific (p = 0.0342) but this was not evident after administration of TCS1102 (p = 0.5108) (Figure 4B). The preference for stressed conspecific was calculated as percentage (Figure 5C); data analysis indicated a significant significant main effect of age (F(1,20) = 5.917; p = 0.0245) and a conspecific age by drug treatment interaction (F(1, 20) = 18.03; p = 0.0004). Orexin receptor inhibition abolished the preference for stressed juveniles (p = 0.0051) and the avoidance of stressed adults (p = 0.0264) (Figure 5C).

**Figure 5.**
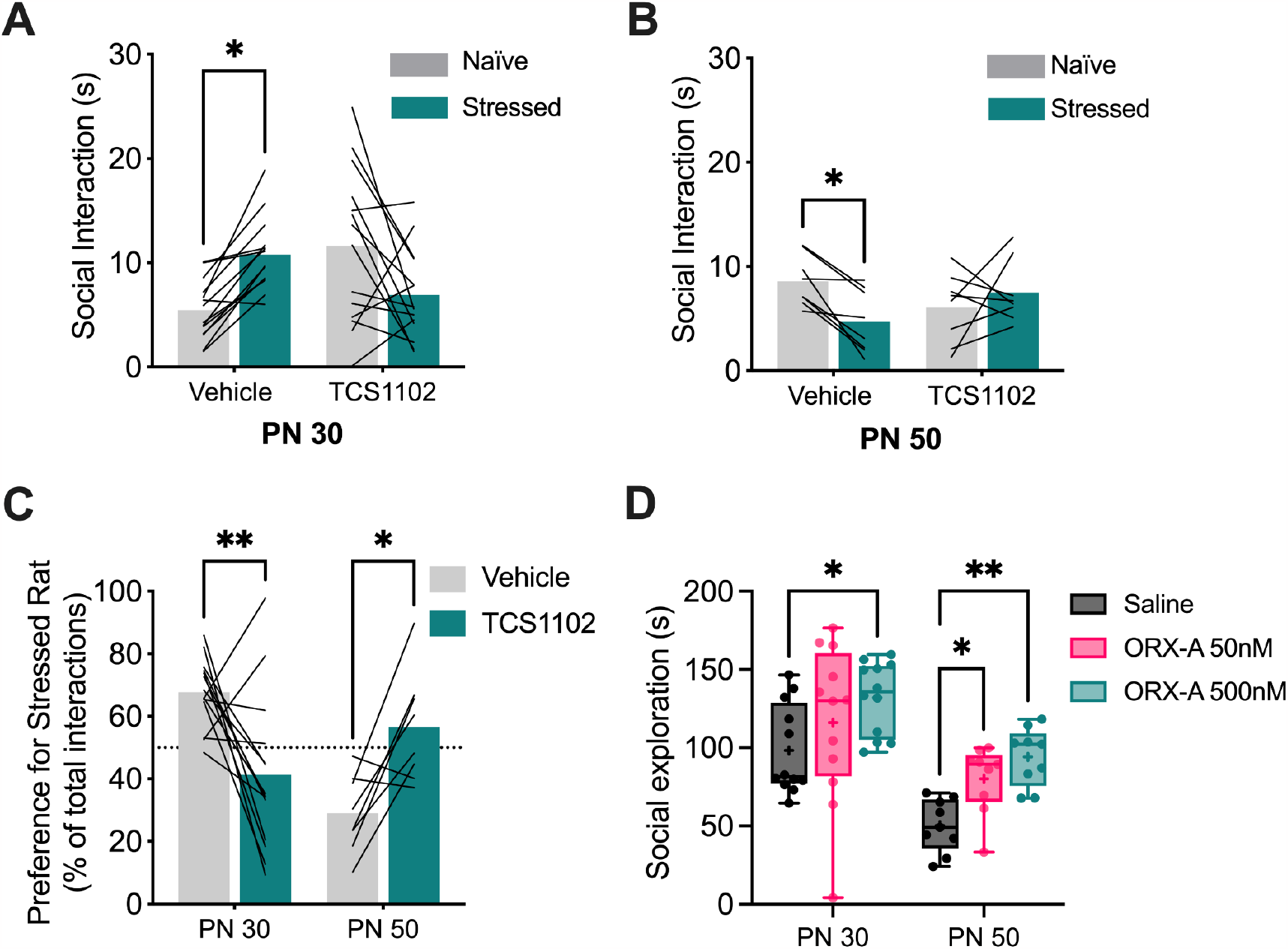
Orexin neurotransmission affects social behavior in rats during SAP and one-on-one social exploration tests when tested with juvenile conspecifics and adult conspecifics. Juvenile and Adult SAP tests were conducted 45 min after administration of the vehicle or the dual orexin receptor antagonist TCS1102 (1µM) to the insula. **A.** Mean (with individual replicates) time spent interacting with juvenile conspecifics during 5 min SAP tests. Test rats spent more time interacting with the stressed juvenile conspecific, this behavior was abolished by TCS1102. **B.** Mean (with individual replicates) time spent interacting with each adult conspecific during 5 min SAP tests. Under vehicle, test rats spent more time interacting with the naïve adult conspecific, which was not evident after administration of TCS1102. **C.** Preference for stressed conspecific in percentage of total interactions time when tested with juvenile or adult conspecifics (Mean with individual replicates). Rats tested with juvenile conspecifics presented a preference for the stressed conspecific, the treatment with TCS1102 significantly reduced this preference. Rats tested with adult conspecifics presented a preference for naïve conspecifics, which was eliminated by insular treatment with TCS1102. **D.** Time spent exploring naïve juvenile and adult conspecifics in a 5 minute social interaction test (box depicts median and first and third quartile, whiskers indicate maximum and minimum, and the mean is indicated by a +) 30 min after insula injections of vehicle or orexin-A (50 and 500 nM). Orexin-A increased time exploring conspecific regardless of age. *p<0.05, **p<0.01.

In the one-on-one social interaction tests, orexin-A appeared to increase interactions toward both naïve juvenile and adult conspecifics (Figure 5D). Data analysis indicated a main effect of conspecific age (F(1, 19) = 22.90; p = 0.0001) and dose (F(2, 38) = 9.906; p = 0.0003). Post-hoc tests indicated that 500nM of orexin-A increased social interaction with naïve juveniles (p = 0.0169) and naïve adults (p = 0.0063).

## DISCUSSION

The current studies investigated how hunger and hunger associated neural circuits and peptides contribute to social decision-making. We demonstrate for the first time that hunger states interfere in the processes of social affective behavior toward conspecifics in distress. Indeed, rats with 14 h of food/water deprivation or relieved with 1 h of food/water access after deprivation preferred to spend more time interacting with naïve juvenile conspecific during SAP test, an opposite behavior from when tested under sated state. Furthermore, in this study we investigated the role of LH and insular MCH and orexin neurotransmission in shaping social affective behavior. Pharmacological inhibition of the LH abolished either preference for stressed juvenile and naïve adult conspecifics. Additionally, we observed that blockade of MCHr1 and orexin receptors in the insula abolished either the preference for stressed juveniles and administration of either MCH or orexin-A increased social interactions directed toward naïve juveniles. Interestingly, only orexin receptors appeared to contribute to behavior in the adult SAP test where blockade of both orexin receptors interfered with social affective behavior while orexin-A itself increased social interaction with naïve adults. Taken together, these findings identify possible mechanisms by which basic motivational states like hunger can interact with socioemotional decisions.

The social decision making process requires the animal to detect and recognize signs of the emotional state of others to generate appropriate behaviors toward each situation, which is important to aggression, mating, maternal behavior and others (Insel and Fernald 2004; Rendon et al. 2015). In this sense, social affective behavior depends on the integration of social affective cues to an appropriate behavior response (Rogers-Carter and Christianson 2019). In addition to external signs of the other individual’s emotional state, this behavioral response also depends on the interoception/internal state (Sandi and Haller 2015; Devlin et al. 2022; Smith and Grueter 2022), which is in line with our findings that the LH is involved in shaping this behavior. Indeed, LH has been related to motivated behavior, stress, arousal, sleep/wake states, feeding behaviors (Bonnavion et al. 2016; Barretto-de-Souza et al. 2022) and, recently, social aspects have also been related to this hypothalamic nucleus (Nieh et al. 2016; Noritake et al. 2020). Indeed, exposing a rat to stress either directly or vicariously also changes social affective preference behaviors (Toyoshima et al. 2021, 2022). Together with our finding that hunger also alters social preference underscores the hypothesis that motivational states interact with social affective processes to organize behavior.

It is tempting to relate the effect of hunger on social behavior to the colloquial experience of “hanger” which describes the unpleasant affect caused by food deprivation. Emotional states have a powerful influence on appraisal of environmental cues and research on humans demonstrated that hunger increased individual’s likelihood to rate social stimuli as aggressive and increased negative social judgements (MacCormack and Lindquist 2019). In the SAP test, we have argued that approach to the stressed juvenile occurs because the juveniles emit socially attractive cues (odors, vocalizations) that are not appraised as threatening, whereas test rats avoid stressed adults because a different set of signals emitted by stressed adults are appraised as social danger signals. In the SAP test, hunger may have caused the test rats to appraise the stressed juvenile as relatively more dangerous leading to affiliation with the unstressed alternative. Prior work on motivational state competition has focused on the AgRP neurons of the arcuate nucleus (Burnett et al. 2016, 2019) which can alter several affective behaviors via connections to the LH (Andermann and Lowell 2017). The literature describes that the downstream connection between the insula and the LH is related to cardiovascular control, feeding and appetitive behaviors (Yasui et al. 1991; Oppenheimer et al. 1992; Wu et al. 2020). However, the upstream pathway, LH to insula, is less studied. Anatomical studies show LH projections in the insula (Kampe et al. 2009; Gehrlach et al. 2020) as well the presence of MCH and orexin peptides (Saito et al. 2001; Jacobson et al. 2022). The current findings that the LH itself and that MCH and orexin acting in the insula influence social affective behavior suggests a tripartite circuit where hunger signals originating in the arcuate nucleus are relayed, at least in part, to the insula via orexin and MCH projection neurons (Kampe et al. 2009).

A central idea relating to motivational state competition is that some needs should outweigh others, such as the need for food prevailing over the need for social interaction (Sutton and Krashes 2020). While this state competition likely unfolds across several levels of neural processing, the insula is site of multisensory integration receiving information from different parts of the brain containing information about social, interoceptive, affective, and motivational states (Krushel and van der Kooy 1988; Paulus and Stein 2006; Damasio and Carvalho 2013; Gogolla et al. 2014; Rogers-Carter and Christianson 2019). Thus, the insula is likely to be central to resolving motivational state conflicts or, more simply, a neural locus for internal states to influence the appraisal of external cues. While we assume this is a part of the cognition underlying partner choice in the SAP test, a similar phenomenon is well described for hunger and thirst. Specifically, insula responses to cues that predict the availability of food and water are dynamically shaped by satiety state (Livneh et al. 2017). A goal of future work will be to determine if insula responses to social affective cues are similarly influenced by hunger, stress and other states that affect the direction of social behavior.

The orexin system is composed of 2 metabotropic receptors called OXR1 and OXR2, and 2 peptide endogenous ligands called orexin-A and orexin-B. The endogenous ligands have different affinity profiles between receptors, orexin-A has high affinity for both OXR1 and OXR2 receptors, while orexin-B has a higher affinity for OXR2 (Jacobson et al. 2022). Orexin neurons are predominantly found in the LH (Swanson et al. 2005). As previously mentioned, we observed that insular treatment with a dual orexin receptor antagonist abolished the preference for the stressed juvenile and naïve adult conspecifics during the SAP test. Also, insula treatment with the endogenous agonist orexin-A increased social exploration time with either naïve juveniles and adults conspecific during the SE test. These results corroborate the fact that the insula would be receiving signals from the internal state and together with social affective information influencing approach/avoidance behavior during SAP test. In this sense, it is known that orexin neurotransmission acts as an important neuropeptide in the process of arousal, stress, motivated behavior and reward, which are important components for social affective behavior (Jacobson et al. 2022). In fact, studies show that orexin stimulation of the posterior insula region promotes an increase in the “liking” reaction to sucrose, showing a hedonic role of this neurochemical mechanism in the insula (Castro and Berridge 2017). It is possible that orexin may contribute to the selection of appetitive or avoidant outputs of the insula via modulation of insular synaptic transmission and integration. Interestingly, OXR1 receptor agonists potentiating GABA mediated fast inhibitory transmission in the insula (Usui et al. 2019) which is necessary for some aspects of typical rodent sociality (Gogolla et al. 2014). The current results warrant further investigation of the cellular distribution of orexin receptors and their postsynaptic signaling mechanisms.

MCH cell populations are also predominantly found in LH (Hahn 2010). The MCH system consists of two class I G-protein-coupled receptors called MCHr1 and MCHr2, and one endogenous ligand, MCH. The MCH system has an important role in feeding behavior and energy balance (Ludwig et al. 2001; Karlsson et al. 2012), but it is also involved in other functions such as sleep/wake (Monti et al. 2013), maternal behaviors (Alachkar et al. 2016) and locomotion (Marsh et al. 2002). Expressions of MCH receptors are different between species, the MCHr1 is widely expressed in rodents and primates brains and the MCHr2 were identified in humans, fishes, dogs, ferrets and non-human primates, but not rodents (Tan et al. 2002; Presse et al. 2014). Thus, the effects observed with the MCH antagonist and agonist should be attributed to MCHr1. The results obtained in this study show that MCH neurotransmission in the insula also influences social affective behavior during SAP tests, however, unlike orexin neurotransmission, the effect of the agonism of MCH receptors was observed only when tested with juvenile conspecifics during SE test. Orexin and MCH neuron populations are found in different locations in the LH and they do not overlap (Hahn 2010), indicating that they may be acting independently. In the insula, MCH appears necessary and sufficient for appetitive social approach behavior to juveniles and so it is possible that both MCH and orexin are released during juvenile social interactions, but only orexin is released in adult interactions. Further, states like hunger may alter the tone of these inputs to the insula as a mechanism for motivational states to interact with social decision making; direct observation of these peptide dynamics will be necessary to test this hypothesis.

In humans, some psychiatric conditions such as autism and schizophrenia present an impairment in detecting the emotional state of others and a subsequent inability to generate appropriate social behaviors (Edwards et al. 2002; Lozier et al. 2014), therefore it is important to better understand the neurobiology of this behavior. Importantly, people with autism spectrum disorder, which is characterized by abnormal communication, social interaction, and impairments in interoception (Quadt et al. 2018), have a decreased volume of LH (Kurth et al. 2011) and numerous differences in insula function (Di Martino et al. 2009). Furthermore, the MCH system is involved in the neurodevelopmental processes, and the deletion of MCHr1 generates schizophrenia-like phenotypes (Vawter et al. 2020). Therefore, these new observations may help identify previously underappreciated targets for therapeutic intervention and new understanding of the pathobiology underlying social and affective symptoms of psychopathology.

## Acknowledgements

The authors wish to thank Nancy McGilloway and Todd Gaines, administrators of the Boston College Animal Care Facility, for outstanding animal husbandry and Bret Judson, director of the Boston College Imaging Core, for training and assistance with microscopy. Funding for this work was provided by the Boston College Undergraduate Research Fellowship, the Boston College McNair Scholars Program, São Paulo Research Foundation (FAPESP) grant #2021/11211-9, and the National Institutes of Health Grant MH119422.

## Notes

**Financial Disclosures:** The authors declare no direct or indirect biomedical financial interests or other potential conflicts of interest.

### Competing Interest Statement

The authors have declared no competing interest.

## REFERENCES

Abbas MdG, Shoji H, Soya S, et al (2015) Comprehensive Behavioral Analysis of Male Ox1r−/− Mice Showed Implication of Orexin Receptor-1 in Mood, Anxiety, and Social Behavior. Front Behav Neurosci 9:

Alachkar A, Alhassen L, Wang Z, et al (2016) Inactivation of the melanin concentrating hormone system impairs maternal behavior. Eur Neuropsychopharmacol J Eur Coll Neuropsychopharmacol 26:1826–1835. https://doi.org/10.1016/j.euroneuro.2016.08.014

Andermann ML, Lowell BB (2017) Toward a Wiring Diagram Understanding of Appetite Control. Neuron 95:757–778. https://doi.org/10.1016/j.neuron.2017.06.014

Barretto-de-Souza L, Benini R, Reis-Silva LL, Crestani CC (2022) Role of CRF1 and CRF2 receptors in the lateral hypothalamus in cardiovascular and anxiogenic responses evoked by restraint stress in rats: Evaluation of acute and chronic exposure. Neuropharmacology 212:109061. https://doi.org/10.1016/j.neuropharm.2022.109061

Blouin AM, Fried I, Wilson CL, et al (2013) Human hypocretin and melanin-concentrating hormone levels are linked to emotion and social interaction. Nat Commun 4:1547. https://doi.org/10.1038/ncomms2461

Bonnavion P, de Lecea L (2010) Hypocretins in the Control of Sleep and Wakefulness. Curr Neurol Neurosci Rep 10:174–179. https://doi.org/10.1007/s11910-010-0101-y

Bonnavion P, Mickelsen LE, Fujita A, et al (2016) Hubs and spokes of the lateral hypothalamus: cell types, circuits and behaviour. J Physiol 594:6443–6462. https://doi.org/10.1113/JP271946

Burnett CJ, Funderburk SC, Navarrete J, et al (2019) Need-based prioritization of behavior. eLife 8:e44527. https://doi.org/10.7554/eLife.44527

Burnett CJ, Li C, Webber E, et al (2016) Hunger-Driven Motivational State Competition. Neuron 92:187–201. https://doi.org/10.1016/j.neuron.2016.08.032

Castro DC, Berridge KC (2017) Opioid and orexin hedonic hotspots in rat orbitofrontal cortex and insula. Proc Natl Acad Sci 114:E9125–E9134. https://doi.org/10.1073/pnas.1705753114

Damasio A, Carvalho GB (2013) The nature of feelings: evolutionary and neurobiological origins. Nat Rev Neurosci 14:143–152. https://doi.org/10.1038/nrn3403

Devlin BA, Smith CJ, Bilbo SD (2022) Sickness and the Social Brain: How the Immune System Regulates Behavior across Species. Brain Behav Evol 97:197–210. https://doi.org/10.1159/000521476

Di Martino A, Ross K, Uddin LQ, et al (2009) Functional Brain Correlates of Social and Nonsocial Processes in Autism Spectrum Disorders: An Activation Likelihood Estimation Meta-Analysis. Biol Psychiatry 65:63–74. https://doi.org/10.1016/j.biopsych.2008.09.022

Djerdjaj A, Ng AJ, Rieger NS, Christianson JP (2022) The basolateral amygdala to posterior insular cortex tract is necessary for social interaction with stressed juvenile rats. Behav Brain Res 435:114050. https://doi.org/10.1016/j.bbr.2022.114050

Edwards J, Jackson HJ, Pattison PE (2002) Emotion recognition via facial expression and affective prosody in schizophrenia: a methodological review. Clin Psychol Rev 22:789–832. https://doi.org/10.1016/s0272-7358(02)00130-7

Gehrlach DA, Gaitanos TN, Klein AS, et al (2020) A whole-brain connectivity map of mouse insular cortex. bioRxiv 2020.02.10.941518. https://doi.org/10.1101/2020.02.10.941518

Gogolla N (2017) The insular cortex. Curr Biol CB 27:R580–R586. https://doi.org/10.1016/j.cub.2017.05.010

Gogolla N, Takesian AE, Feng G, et al (2014) Sensory Integration in Mouse Insular Cortex Reflects GABA Circuit Maturation. Neuron 83:894–905. https://doi.org/10.1016/j.neuron.2014.06.033

Hagar JM, Macht VA, Wilson SP, Fadel JR (2017) Upregulation of orexin/hypocretin expression in aged rats: Effects on feeding latency and neurotransmission in the insular cortex. Neuroscience 350:124–132. https://doi.org/10.1016/j.neuroscience.2017.03.021

Hahn JD (2010) Comparison of melanin-concentrating hormone and hypocretin/orexin peptide expression patterns in a current parceling scheme of the lateral hypothalamic zone. Neurosci Lett 468:12–17. https://doi.org/10.1016/j.neulet.2009.10.047

Hervieu GJ, Cluderay JE, Harrison D, et al (2000) The distribution of the mRNA and protein products of the melanin-concentrating hormone (MCH) receptor gene, slc-1, in the central nervous system of the rat. Eur J Neurosci 12:1194–1216. https://doi.org/10.1046/j.1460-9568.2000.00008.x

Hsiao Y-T, Jou S-B, Yi P-L, Chang F-C (2012) Activation of GABAergic pathway by hypocretin in the median raphe nucleus (MRN) mediates stress-induced theta rhythm in rats. Behav Brain Res 233:224–231. https://doi.org/10.1016/j.bbr.2012.05.002

Huang H, Acuna-Goycolea C, Li Y, et al (2007) Cannabinoids excite hypothalamic melanin-concentrating hormone but inhibit hypocretin/orexin neurons: implications for cannabinoid actions on food intake and cognitive arousal. J Neurosci Off J Soc Neurosci 27:4870–4881. https://doi.org/10.1523/JNEUROSCI.0732-07.2007

Insel TR, Fernald RD (2004) HOW THE BRAIN PROCESSES SOCIAL INFORMATION: Searching for the Social Brain. Annu Rev Neurosci 27:697–722. https://doi.org/10.1146/annurev.neuro.27.070203.144148

Jacobson LH, Hoyer D, de Lecea L (2022) Hypocretins (orexins): The ultimate translational neuropeptides. J Intern Med 291:533–556. https://doi.org/10.1111/joim.13406

Kampe J, Tschöp MH, Hollis JH, Oldfield BJ (2009) An anatomic basis for the communication of hypothalamic, cortical and mesolimbic circuitry in the regulation of energy balance. Eur J Neurosci 30:415–430. https://doi.org/10.1111/j.1460-9568.2009.06818.x

Kaplan GB, Lakis GA, Zhoba H (2022) Sleep-wake and arousal dysfunctions in post-traumatic stress disorder: Role of orexin systems. Brain Res Bull 186:106–122. https://doi.org/10.1016/j.brainresbull.2022.05.006

Karlsson C, Zook M, Ciccocioppo R, et al (2012) Melanin-concentrating hormone receptor 1 (MCH1-R) antagonism: Reduced appetite for calories and suppression of addictive-like behaviors. Pharmacol Biochem Behav 102:400–406. https://doi.org/10.1016/j.pbb.2012.06.010

Kokare DM, Patole AM, Carta A, et al (2006) GABAA receptors mediate orexin-A induced stimulation of food intake. Neuropharmacology 50:16–24. https://doi.org/10.1016/j.neuropharm.2005.07.019

Krushel LA, van der Kooy D (1988) Visceral cortex: integration of the mucosal senses with limbic information in the rat agranular insular cortex. J Comp Neurol 270:39–54, 62–63. https://doi.org/10.1002/cne.902700105

Kurth F, Narr KL, Woods RP, et al (2011) Diminished Gray Matter Within the Hypothalamus in Autism Disorder: A Potential Link to Hormonal Effects? Biol Psychiatry 70:278–282. https://doi.org/10.1016/j.biopsych.2011.03.026

Kurth F, Zilles K, Fox PT, et al (2010) A link between the systems: functional differentiation and integration within the human insula revealed by meta-analysis. Brain Struct Funct 214:519–534. https://doi.org/10.1007/s00429-010-0255-z

Li S-B, de Lecea L (2020) The hypocretin (orexin) system: from a neural circuitry perspective. Neuropharmacology 167:107993. https://doi.org/10.1016/j.neuropharm.2020.107993

Li T-L, Lee Y-H, Wu F-H, Hwang L-L (2021) Orexin-A directly depolarizes dorsomedial hypothalamic neurons, including those innervating the rostral ventrolateral medulla. Eur J Pharmacol 899:174033. https://doi.org/10.1016/j.ejphar.2021.174033

Liu J-J, Tsien RW, Pang ZP (2022) Hypothalamic melanin-concentrating hormone regulates hippocampus-dorsolateral septum activity. Nat Neurosci 25:61–71. https://doi.org/10.1038/s41593-021-00984-5

Livneh Y, Andermann ML (2021) Cellular activity in insular cortex across seconds to hours: Sensations and predictions of bodily states. Neuron 109:3576–3593. https://doi.org/10.1016/j.neuron.2021.08.036

Livneh Y, Ramesh RN, Burgess CR, et al (2017) Homeostatic circuits selectively gate food cue responses in insular cortex. Nature 546:611–616. https://doi.org/10.1038/nature22375

Lozier LM, Vanmeter JW, Marsh AA (2014) Impairments in facial affect recognition associated with autism spectrum disorders: A meta-analysis. Dev Psychopathol 26:933–945. https://doi.org/10.1017/S0954579414000479

Ludwig DS, Tritos NA, Mastaitis JW, et al (2001) Melanin-concentrating hormone overexpression in transgenic mice leads to obesity and insulin resistance. J Clin Invest 107:379–386. https://doi.org/10.1172/JCI10660

MacCormack JK, Lindquist KA (2019) Feeling hangry? When hunger is conceptualized as emotion. Emotion 19:301–319. https://doi.org/10.1037/emo0000422

Marsh DJ, Weingarth DT, Novi DE, et al (2002) Melanin-concentrating hormone 1 receptor-deficient mice are lean, hyperactive, and hyperphagic and have altered metabolism. Proc Natl Acad Sci 99:3240–3245. https://doi.org/10.1073/pnas.052706899

Matsuki T, Nomiyama M, Takahira H, et al (2009) Selective loss of GABAB receptors in orexin-producing neurons results in disrupted sleep/wakefulness architecture. Proc Natl Acad Sci 106:4459–4464. https://doi.org/10.1073/pnas.0811126106

Meyza KZ, Bartal IB-A, Monfils MH, et al (2017) The roots of empathy: Through the lens of rodent models. Neurosci Biobehav Rev 76:216–234. https://doi.org/10.1016/j.neubiorev.2016.10.028

Mitsukawa K, Kimura H (2022) Orexin 2 receptor (OX2R) protein distribution measured by autoradiography using radiolabeled OX2R-selective antagonist EMPA in rodent brain and peripheral tissues. Sci Rep 12:8473. https://doi.org/10.1038/s41598-022-12601-x

Monti JM, Torterolo P, Lagos P (2013) Melanin-concentrating hormone control of sleep-wake behavior. Sleep Med Rev 17:293–298. https://doi.org/10.1016/j.smrv.2012.10.002

Newman SW (1999) The Medial Extended Amygdala in Male Reproductive Behavior A Node in the Mammalian Social Behavior Network. Ann N Y Acad Sci 877:242–257. https://doi.org/10.1111/j.1749-6632.1999.tb09271.x

Nieh EH, Vander Weele CM, Matthews GA, et al (2016) Inhibitory Input from the Lateral Hypothalamus to the Ventral Tegmental Area Disinhibits Dopamine Neurons and Promotes Behavioral Activation. Neuron 90:1286–1298. https://doi.org/10.1016/j.neuron.2016.04.035

Noritake A, Ninomiya T, Isoda M (2020) Representation of distinct reward variables for self and other in primate lateral hypothalamus. Proc Natl Acad Sci 117:5516–5524. https://doi.org/10.1073/pnas.1917156117

O’Connell LA, Hofmann HA (2011) The Vertebrate mesolimbic reward system and social behavior network: A comparative synthesis. J Comp Neurol 519:3599–3639. https://doi.org/10.1002/cne.22735

Oppenheimer SM, Saleh T, Cechetto DF (1992) Lateral hypothalamic area neurotransmission and neuromodulation of the specific cardiac effects of insular cortex stimulation. Brain Res 581:133–142. https://doi.org/10.1016/0006-8993(92)90352-A

Paulus MP, Stein MB (2006) An insular view of anxiety. Biol Psychiatry 60:383–387. https://doi.org/10.1016/j.biopsych.2006.03.042

Petrovich GD (2018) Lateral Hypothalamus as a Motivation-Cognition Interface in the Control of Feeding Behavior. Front Syst Neurosci 12:

Presse F, Conductier G, Rovere C, Nahon J-L (2014) The melanin-concentrating hormone receptors: neuronal and non-neuronal functions. Int J Obes Suppl 4:S31–S36. https://doi.org/10.1038/ijosup.2014.9

Quadt L, Critchley HD, Garfinkel SN (2018) The neurobiology of interoception in health and disease. Ann N Y Acad Sci 1428:112–128. https://doi.org/10.1111/nyas.13915

Ranote S, Elliott R, Abel KM, et al (2004) The neural basis of maternal responsiveness to infants: an fMRI study. NeuroReport 15:1825. https://doi.org/10.1097/01.wnr.0000137078.64128.6a

Rendon NM, Keesom SM, Amadi C, et al (2015) Vocalizations convey sex, seasonal phenotype, and aggression in a seasonal mammal. Physiol Behav 152:143–150. https://doi.org/10.1016/j.physbeh.2015.09.014

Rieger NS, Varela JA, Ng AJ, et al (2022) Insular cortex corticotropin-releasing factor integrates stress signaling with social affective behavior. Neuropsychopharmacology 47:1156–1168. https://doi.org/10.1038/s41386-022-01292-7

Rodrigues LTC, Patrone LGA, Gargaglioni LH, Dias MB (2022) Melanin-concentrating hormone regulates the hypercapnic chemoreflex by acting in the lateral hypothalamic area. Exp Physiol 107:1298–1311. https://doi.org/10.1113/EP090318

Rogers-Carter MM, Christianson JP (2019) An insular view of the social decision-making network. Neurosci Biobehav Rev 103:119–132. https://doi.org/10.1016/j.neubiorev.2019.06.005

Rogers-Carter MM, Djerdjaj A, Gribbons KB, et al (2019) Insular Cortex Projections to Nucleus Accumbens Core Mediate Social Approach to Stressed Juvenile Rats. J Neurosci Off J Soc Neurosci 39:8717–8729. https://doi.org/10.1523/JNEUROSCI.0316-19.2019

Rogers-Carter MM, Varela JA, Gribbons KB, et al (2018) Insular cortex mediates approach and avoidance responses to social affective stimuli. Nat Neurosci 21:404–414. https://doi.org/10.1038/s41593-018-0071-y

Sabetghadam A, Grabowiecka-Nowak A, Kania A, et al (2018) Melanin-concentrating hormone and orexin systems in rat nucleus incertus: Dual innervation, bidirectional effects on neuron activity, and differential influences on arousal and feeding. Neuropharmacology 139:238–256. https://doi.org/10.1016/j.neuropharm.2018.07.004

Saito Y, Cheng M, Leslie FM, Civelli O (2001) Expression of the melanin-concentrating hormone (MCH) receptor mRNA in the rat brain. J Comp Neurol 435:26–40. https://doi.org/10.1002/cne.1191

Sakurai T, Amemiya A, Ishii M, et al (1998) Orexins and orexin receptors: a family of hypothalamic neuropeptides and G protein-coupled receptors that regulate feeding behavior. Cell 92:573–585. https://doi.org/10.1016/s0092-8674(00)80949-6

Sanathara N, Alhassen L, Marmouzi I, et al (2021) Oxytocin-MCH circuit regulates monosynaptic inputs to MCH neurons and modulates social recognition memory. Neuropharmacology 184:108423. https://doi.org/10.1016/j.neuropharm.2020.108423

Sandi C, Haller J (2015) Stress and the social brain: behavioural effects and neurobiological mechanisms. Nat Rev Neurosci 16:290–304. https://doi.org/10.1038/nrn3918

Saper CB, Swanson LW, Cowan WM (1979) An autoradiographic study of the efferent connections of the lateral hypothalamic area in the rat. J Comp Neurol 183:689–706. https://doi.org/10.1002/cne.901830402

Smith NK, Grueter BA (2022) Hunger-driven adaptive prioritization of behavior. FEBS J 289:922–936. https://doi.org/10.1111/febs.15791

Sutton AK, Krashes MJ (2020) Integrating Hunger with Rival Motivations. Trends Endocrinol Metab TEM 31:495–507. https://doi.org/10.1016/j.tem.2020.04.006

Swanson LW, Sanchez-Watts G, Watts AG (2005) Comparison of melanin-concentrating hormone and hypocretin/orexin mRNA expression patterns in a new parceling scheme of the lateral hypothalamic zone. Neurosci Lett 387:80–84. https://doi.org/10.1016/j.neulet.2005.06.066

Tan CP, Sano H, Iwaasa H, et al (2002) Melanin-Concentrating Hormone Receptor Subtypes 1 and 2: Species-Specific Gene Expression. Genomics 79:785–792. https://doi.org/10.1006/geno.2002.6771

Toyoshima M, Mitsui K, Yamada K (2021) Prior stress experience modulates social preference for stressed conspecifics in male rats. Neurosci Lett 765:136253. https://doi.org/10.1016/j.neulet.2021.136253

Toyoshima M, Okuda E, Hasegawa N, et al (2022) Socially Transferred Stress Experience Modulates Social Affective Behaviors in Rats. Neuroscience 502:68–76. https://doi.org/10.1016/j.neuroscience.2022.08.022

Usui M, Kaneko K, Oi Y, Kobayashi M (2019) Orexin facilitates GABAergic IPSCs via postsynaptic OX1 receptors coupling to the intracellular PKC signalling cascade in the rat cerebral cortex. Neuropharmacology 149:97–112. https://doi.org/10.1016/j.neuropharm.2019.02.012

Vawter MP, Schulmann A, Alhassen L, et al (2020) Melanin Concentrating Hormone Signaling Deficits in Schizophrenia: Association With Memory and Social Impairments and Abnormal Sensorimotor Gating. Int J Neuropsychopharmacol 23:53–65. https://doi.org/10.1093/ijnp/pyz051

Wu Y, Chen C, Chen M, et al (2020) The anterior insular cortex unilaterally controls feeding in response to aversive visceral stimuli in mice. Nat Commun 11:640. https://doi.org/10.1038/s41467-020-14281-5

Yasui Y, Breder CD, Saper CB, Cechetto DF (1991) Autonomic responses and efferent pathways from the insular cortex in the rat. J Comp Neurol 303:355–374. https://doi.org/10.1002/cne.903030303

Yiannakas A, Rosenblum K (2017) The Insula and Taste Learning. Front Mol Neurosci 10:335. https://doi.org/10.3389/fnmol.2017.00335

